# Biased M_1_ positive allosteric modulators reveal novel role of phospholipase D in M_1_-dependent rodent cortical plasticity

**DOI:** 10.1101/806943

**Authors:** Sean P. Moran, Zixiu Xiang, Catherine A. Doyle, James Maksymetz, Xiaohui Lv, Sehr Faltin, Colleen M. Niswender, Jerri M. Rook, Craig W. Lindsley, P. Jeffrey Conn

## Abstract

Highly selective positive allosteric modulators (PAMs) of the M_1_ subtype of muscarinic acetylcholine receptor have emerged as an exciting new approach for the potential improvement of cognitive function in patients suffering from Alzheimer’s disease and schizophrenia. M_1_ PAM discovery programs have produced a structurally diverse range of M_1_ PAMs with distinct pharmacological properties, including different levels of agonist activity and differences in signal bias. This includes the recent discovery of novel biased M_1_ PAMs that can potentiate coupling of M_1_ to activation of phospholipase C but not phospholipase D (PLD). However, little is known about the role of PLD in M_1_ signaling in native systems and it is not clear whether biased M_1_ PAMs will display differences in modulating M_1_-mediated responses in native tissue. We now report a series of studies using novel PLD inhibitors and PLD knockout mice to show that PLD is necessary for the induction of M_1_-dependent long-term depression (LTD) in the prefrontal cortex (PFC). Importantly, biased M_1_ PAMs that do not couple to PLD not only fail to potentiate orthosteric agonist-induced LTD but also block M_1_-dependent LTD in the PFC. In contrast, biased and non-biased M_1_ PAMs act similarly in potentiating M_1_-dependent electrophysiological responses that are PLD-independent. These findings demonstrate that PLD plays a critical role in the ability of M_1_ PAMs to modulate certain CNS functions and that biased M_1_ PAMs function differently in brain regions implicated in cognition.

**Summary:** We demonstrate a novel role of phospholipase D in M_1_-dependent rodent cortical plasticity and M_1_ PAMs that do not couple to phospholipase D have functionally distinct effects on cortical plasticity than non-biased M_1_ PAMs.

## Introduction

The M_1_ muscarinic acetylcholine receptor (mAChR) has attracted intense interest as a promising therapeutic target for the treatment of the cognitive disruptions in schizophrenia and Alzheimer’s disease (AD). M_1_ is highly expressed across many forebrain regions implicated in the pathophysiology of schizophrenia and AD, including the cortex, striatum, and hippocampus (*1, 2*). Interestingly, dysregulation of the M_1_ receptor has been reported within a subset of patients suffering from schizophrenia illustrated by a dramatic reduction of M_1_ receptor expression in pyramidal neurons in cortical regions highly implicated in complex behaviors such as cognition and working memory (*3, 4*). Furthermore, cholinesterase inhibitors, which boost overall cholinergic tone, provide some efficacy in patients with AD; however, dose-limiting adverse effects typically occur with disease progression. Therefore, selectively enhancing M_1_ signaling may provide a potential therapeutic approach for the treatment of the cognitive deficits associated with AD and schizophrenia.

Several orthosteric mAChR agonists, including the nonselective mAChR partial agonist xanomeline (*5*), have entered clinical trials as potential cognition-enhancing agents. Unfortunately, xanomeline failed to meet significant cognition enhancement end points, a result attributed to dose-limiting nonselective cholinergic agonist adverse effects hypothesized to be mediated by the activation of peripheral M_2_ and M_3_ receptors (*6–9*). To increase selectivity for M_1_ and therefore minimize nonselective adverse effects, multiple research efforts shifted to developing compounds that act via allosteric sites on mAChRs, which are structurally distinct from the orthosteric binding site and may be less highly conserved among receptor subtypes. To date, we and others have identified highly subtype-selective positive allosteric modulators (PAMs) of the M_1_ receptor that avoid activation of other mAChR subtypes (see reviews (*10–12*)). Importantly, M_1_ PAMs have shown robust efficacy in enhancing cognition and rescuing cognitive deficits in preclinical animal models relevant for AD and schizophrenia (*13–18*).

While these preclinical findings are extremely promising for the potential of M_1_ PAMs to reverse cognitive deficits in patients, M_1_ PAMs can display a diverse range of pharmacological properties, some of which are potentially detrimental to in vivo efficacy. Previously, we found that the presence of allosteric agonist activity in M_1_ PAMs can limit in vivo efficacy and increase adverse effect liability (*13, 14, 19, 20*). Thus, minimalizing agonist activity can maximize the therapeutic window of M_1_ PAMs (*13, 19–21*). These previous studies demonstrate that a complete understanding of the different pharmacological properties of structurally distinct M_1_ PAMs is essential to fully evaluate clinical candidates and maximize their therapeutic potential.

In addition to displaying differences in allosteric agonist versus pure PAM activity, M_1_ PAMs can also differ in their ability to confer bias to M_1_ signaling. Signal bias is the phenomenon in which different GPCR ligands induce unique active receptor-complex states that are biased toward specific signaling pathways (*22*). For instance, characterization of a broad range of structurally diverse M_1_ PAMs revealed that some M_1_ PAMs potentiate M_1_ signaling through the canonical phospholipase C (PLC) pathway but do not potentiate M_1_-mediated activation of another phospholipase, phospholipase D (PLD) (*23*). PLD is a widely expressed enzyme that hydrolyzes the major membrane phospholipid phosphatidylcholine into the signaling molecules phosphatidic acid (PA) and choline (*24*). PLD can be activated by a variety of receptors, including the M_1_ receptor (*25, 26*). While there are six distinct mammalian isoforms of PLD, only PLD_1_ and PLD_2_ have well established enzymatic activity within the CNS (*24, 27*). However, little is known about the roles of PLD in regulating brain function, and the potential roles of PLD in M_1_-dependent signaling has not been explored. Thus, it is unclear whether M_1_ PAMs that do not activate coupling of M_1_ to PLD in cell lines will display functional differences in regulating M_1_ signaling in the CNS relative to non-biased M_1_ PAMs. For other GPCRs, signal bias provides the exciting potential advantage of selectively activating or potentiating therapeutically relevant pathways while minimizing activation of pathways responsible for adverse effects (*11, 28, 29*). Therefore, a better understanding of these signaling mechanisms is essential for the development of M_1_ PAMs as potential therapeutics for the treatment of prevalent cognitive disorders.

We now report that PLD activity is necessary for a form of M_1_-dependent long-term depression (LTD) in the prefrontal cortex (PFC) that has previously been implicated in the potential therapeutic response to M_1_ PAMs (*14, 30, 31*). Furthermore, biased M_1_ PAMs that do not potentiate M_1_ coupling to PLD fail to potentiate this form of LTD, but actively inhibit M_1_-dependent LTD at this synapse. In contrast, biased and non-biased M_1_ PAMs function similarly in their ability to potentiate M_1_-dependent responses in the CNS that we found to be PLD-independent. Together, these studies reveal that PLD is a critical downstream signaling node for this M_1_-dependent LTD in the PFC and demonstrate that biased M_1_ PAMs can have fundamentally different effects, relative to those of non-biased M_1_ PAMs, in regulating specific aspects of CNS function.

## Results

### M_1_ activation leads to phospholipase D activity in hM_1_-CHO cells

M_1_ activation has previously been shown to lead to an increase in PLD activity (*25, 26*), but it is not known whether this reflects activation of PLD_1_, PLD_2_, or both isoforms. Therefore, we set out to characterize the relative contribution of the two distinct PLD isoforms on selective activation of M_1_ in Chinese Hamster Ovary (CHO) cells stably expressing the M_1_ receptor. While direct measurement of the PLD product PA is challenging due to its rapid conversion into other lipids such as diacylglycerol and lysophosphatidic acid, in the presence of a primary alcohol such as 1-butanol, PLD generates the stable product phosphatidylbutanol (pButanol). pButanol cannot be metabolized, therefore allowing for quantification of intracellular PLD activity (*24, 32, 33*). In agreement with previous findings (*23*), the cholinergic orthosteric agonist carbachol (CCh) induces a robust increase in pButanol accumulation that is blocked by the selective M_1_ antagonist VU0255035 (*34*) (Fig. 1A). Furthermore, the PLD_1,2_ inhibitor ML299 (*35*) potently blocks M_1_-mediated pButanol production, thereby validating that PLD is responsible for the generation of pButanol. Using the recently developed PLD_1_ (VU0359595) (*36*) and PLD_2_ (VU0364739) (*37*) isoform-selective inhibitors, we report that pharmacological inhibition of PLD_1_, but not PLD_2_, blocks M_1_-dependent activation of PLD in this in vitro assay (Fig. 1A). These data reveal that in this cell-based assay, M_1_-dependent activation of PLD primarily occurs through PLD_1_, not PLD_2_.

**Figure 1:**
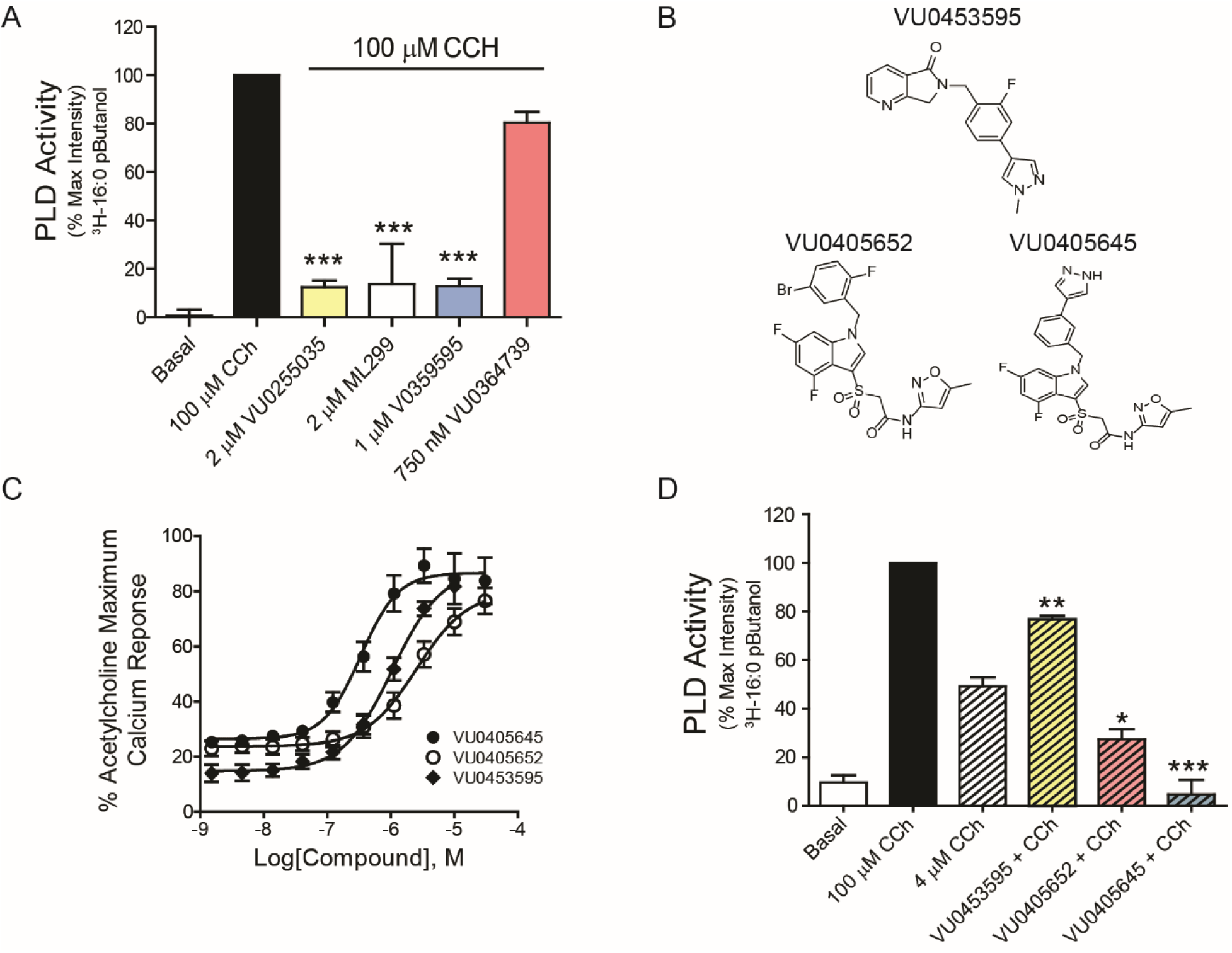
M_1_ activation leads to phospholipase D (PLD) activity in hM_1_-CHO cells and M1 PAMs show differential signal bias in potentiating M_1_–mediated PLD signaling. (A) In rM_1_ CHO cells, PLD activity is measured by quantification of the PLD product phosphatidylbutanol (pButanol) normalized to 100 µM CCh. PLD activity is blocked by 2 µM VU0255035 (M_1_ antagonist, p < 0.001 relative to 100 µM CCh), 2 µM ML299 (PLD_1,2_ inhibitor, p < 0.001 relative to 100 µM CCh), 1 µM VU0359595 (PLD_1_ inhibitor, p < 0.001 relative to 100 µM CCh), but not by 750 nM VU0364739 (PLD_2_ inhibitor, p > 0.05 relative to 100 µM CCh) (one-way ANOVA F_4,10_ = 29.34; p = 0.0001, with a post-hoc Dunnett’s test using CCh alone as the control group, *** p < 0.001). B) Structures of the M_1_ PAMs VU0453595 (left), VU0405652 (center), VU0405645 (right). (C) Concentration−response curves of rM_1_-CHO calcium mobilization assay for VU0453595, VU0405652, VU0405645 in the presence of an EC_20_ concentration of acetylcholine. (D) 4 µM CCh produces a submaximal increase in PLD activity that can be potentiated by 10 µM VU0453595 (p < 0.01 relative to 4 µM CCh). 10 µM VU0405652 (p < 0.05 relative to 4 µM CCh) and 10 µM VU0405645 (p < 0.001 relative to 4 µM CCh) significantly reduced PLD activity compared to 4 µM CCh alone (one-way ANOVA F_3,8_ = 55.1; p = 0.0001, *p < 0.05, **p < 0.01, *** p < 0.001). Data from (A), (C) and (D) represent mean ± SEM from three independent experiments performed in triplicate.

While we have previously characterized M_1_ PAMs that couple to PLC but not PLD (*23*), these early biased M_1_ PAMs suffer from low potency and aqueous solubility. Therefore, we set out to develop additional M_1_ PAMs that potentiate M_1_ coupling to PLC, but do not potentiate coupling to PLD. Previously, we have reported the M_1_ PAMs VU0453595 (half-maximal effective concentration (EC_50_) 2140 ± 440 nM), VU0405652 (EC_50_ 2580 ± 440 nM) and VU0405645 (EC_50_ 340 ± 30 nM) are potent M_1_ PAMs with respect to their ability to potentiate calcium mobilization in CHO cells stably expressing the M_1_ receptor (Fig. 1B,C) (*30, 38*). We now report, unlike the prototypical M1 PAM VU0453595, that both VU0405652 and VU0405645 fail to potentiate CCh activation of PLD in this in vitro assay (Fig. 1D). These findings demonstrate that VU0405652 and VU0405645 but not VU0453595 are biased M_1_ PAMs in that they do not potentiate M_1_ coupling to PLD.

### PLD_1_ but not PLD_2_ is required for M_1_-mediated LTD in the mPFC

From the cell-based assay we know that M_1_ activation can increase PLD activity; however, little is known about whether PLD is necessary for M_1_-dependent responses in native neuronal tissue. Therefore, we set out to characterize the role of PLD in mediating established responses to M_1_ activation in CNS preparations. One response to M_1_ activation that may be relevant to some aspects of cognitive function is induction of long-term depression (LTD) of excitatory synaptic transmission in the medial prefrontal cortex (mPFC) (*13, 14, 19, 30, 31, 39*). We assessed the role of PLD in inducing LTD in the mPFC by measuring changes in layer V field excitatory postsynaptic potentials (fEPSP) evoked by electrical stimulation of afferents in layer II/III of the mPFC (Fig. 2A). Consistent with previous findings (*13, 14, 30, 39*), a maximal concentration of CCh induces robust LTD of fEPSP slope at this synapse (Fig. 2B). Interestingly, bath application of the PLD_1,2_ inhibitor ML299 before and during CCh application led to a complete loss of CCh-induced LTD (Fig. 2C, F). Using selective inhibitors for each PLD isoform, we found that pharmacological inhibition of PLD_1_ with VU0359595 fully blocked CCh-induced LTD (Fig 2D, F). Congruent with the cell-based assay findings, inhibition of PLD_2_ with VU0364739 had no effect on CCh-induced LTD at this synapse (Fig. 2E, F).

**Figure 2:**
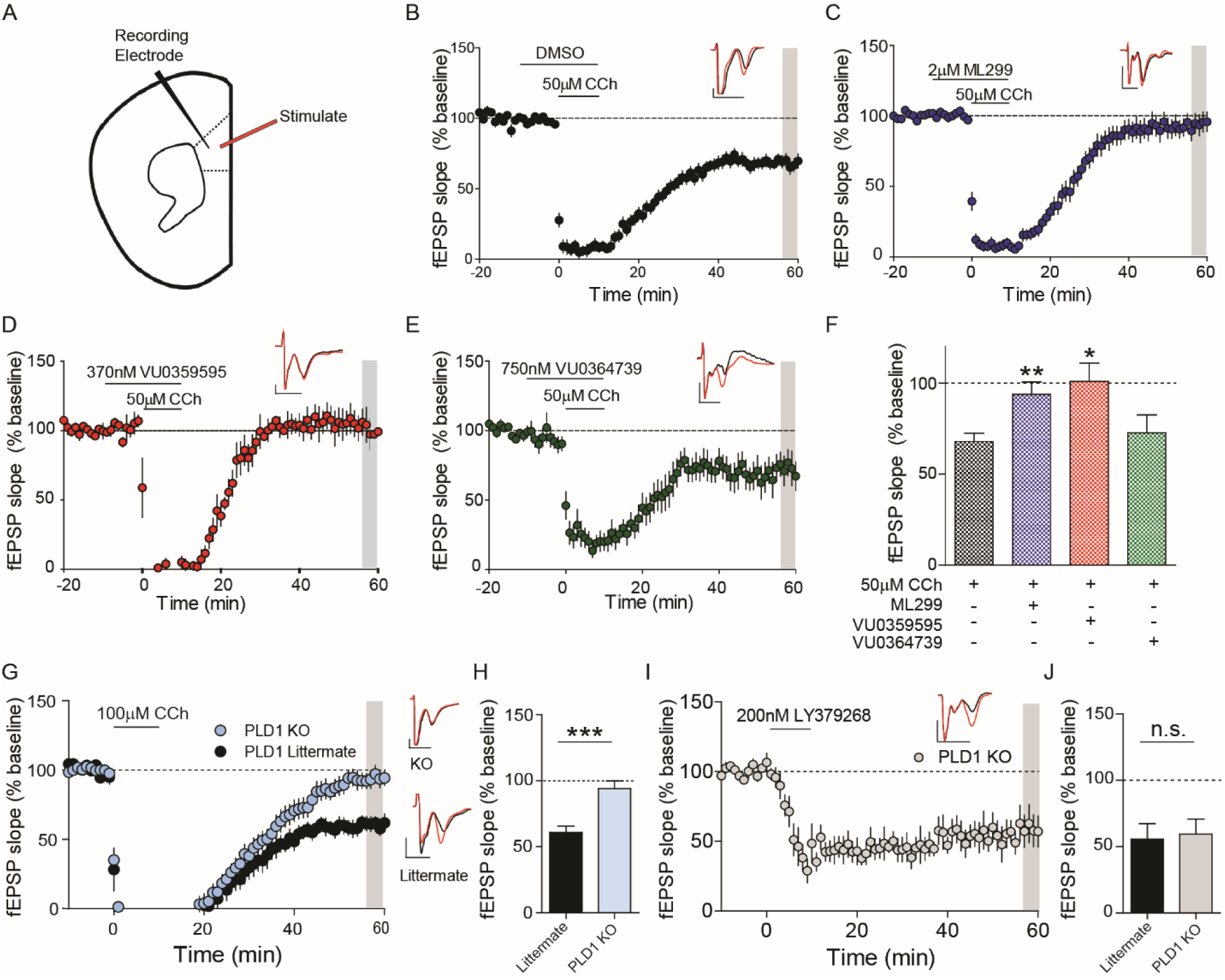
PLD_1_ but not PLD_2_ is necessary for CCh-LTD in the medial prefrontal cortex (mPFC). (A) Schematic of the field excitatory post-synaptic potentials (fEPSP) recorded from layer V of the mouse mPFC in response to electrical stimulation in the superficial layers II–III. (B) Time course graph showing that bath application of 50 μM carbachol (CCh) induces a long-term depression of fEPSPs (68.0 ± 4.44%, n/N= 26/20 slices/mice). (C) 10 min pretreatment of the PLD_1,2_ inhibitor ML299 (2 μM), followed by a 10 min co-application of ML299 and 50 μM CCh, blocked CCh-LTD (93.8 ± 6.74%, n/N= 21/10). (D) Bath application of the PLD_1_-specific inhibitor VU0359595 (370 nM) blocked CCh-LTD (101 ± 10.1%, n/N= 7/6) whereas the PLD_2_ selective inhibitor (E) VU0364739 (750 nM) had no effect on CCh-LTD (69.3 ± 13.0, n/N= 8/4). (F) Quantification of fEPSP slope 46–50 min following drug washout (shaded area) indicates a significant difference between 50 μM CCh alone and ML299, VU0359595 but not VU0364739 (one-way ANOVA F_3,58_ = 5.21; p = 0.0029, with a post-hoc Dunnett’s test using CCh alone as the control group, * p < 0.05, **p < 0.01). (G) Left: Time course graph showing that bath application of 100 μM CCh induces LTD in littermate controls (59.6 ± 6.06 n/N= 9/6) but not PLD_1_ KO mice (92.2 ± 3.21, n/N= 9/6). Right: sample traces for PLD_1_ KO animals (Top) and littermate controls (Bottom). (H) Quantification of fEPSP slope 46–50 min following drug washout (shaded area) indicates a significant difference between genotype (Student’s t-test; p = 0.0002, ***p < 0.001). (I) Bath application of the group II metabotropic glutamate receptor agonist LY379268 (200 nM) for 10 min induces a robust LTD of fEPSP slope in PLD_1_ KO mice (59.4 ± 11.4%, n/N = 5/3). (J) Summary bar graph indicates no significant different between PLD_1_ KO mice and littermate controls (55.7 ± 11.6%, n/N = 7/3; Student’s t-test; p = 0.828). Inset shows representative fEPSP traces for each condition for baseline (red trace) and 50 min after CCh washout (black trace). Scale bars denote 0.25 mV and 5 ms. Data are expressed as mean ± SEM.

To confirm these pharmacological results, we obtained PLD_1_ knockout (KO) mice. In agreement with the pharmacological findings, CCh induces a robust LTD in the littermate controls but not the PLD_1_ KO mice (Fig. 2G, H). Importantly, the ability of a selective agonist of group II metabotropic glutamate receptors (LY379268) to induce LTD was intact in the PLD_1_ KO mice and not significantly different than littermate controls (Fig. I, J). This form of LTD has been previously characterized in detail and is mechanistically distinct from M_1_-dependent LTD in the mPFC (*40–42*). These data suggest that the loss of M_1_-mediated LTD is not due to a general deficit in LTD in this brain region. Taken together, these data demonstrate a critical role of PLD, specifically PLD_1_, in this form of cortical M_1_-LTD.

### Biased M_1_ PAMs fail to potentiate M_1_-LTD in the mPFC

We next tested the hypothesis that biased and non-biased M_1_ PAMs would display functional differences in their ability to potentiate this PLD-dependent CCh-induced LTD of layer V fEPSPs electrically evoked in layer II/III in the mPFC. As previously shown, a submaximal concentration of CCh (10 µM) does not induce LTD at this synapse (Fig. 3A) (*13, 19, 30*). Similar to previous findings, bath application of the non-biased M_1_ PAM VU0453595 for 10 min before and during CCh application leads to a robust LTD (Fig. 3B). Consistent with a role of PLD in inducing M_1_-LTD, neither of the biased M_1_ PAMs, VU0405652 (Fig. 3C) nor VU0405645 (Fig. 3D), potentiate the LTD response to a submaximal concentration of CCh. Quantification of fEPSP slope following drug washout indicates a significant depression of fEPSP slope compared to baseline with the M_1_ PAM VU0453595 but not VU0405652 nor VU0405645 (Fig. 3E).

**Figure 3:**
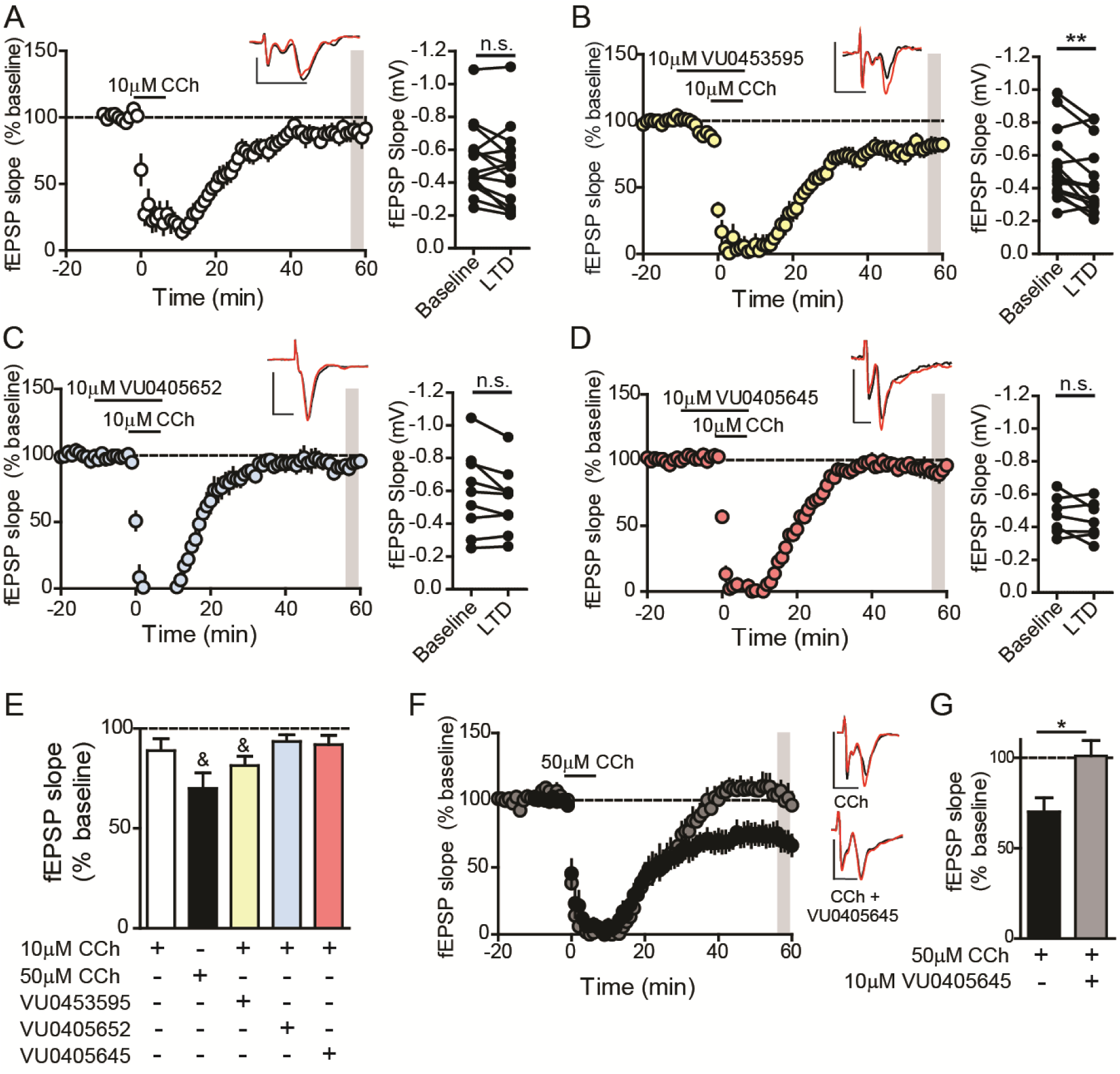
Biased M_1_ PAMs fail to potentiate a submaximal mLTD in the mPFC and actively block CCh-LTD. (A) Time course graph showing that bath application of 10 μM carbachol (CCh) induces a minimal long-term depression (LTD) of fEPSPs (88.9 ± 6.05, n/N= 15/13 slices/mice). Comparison of fEPSPs during baseline and 46-50 min after drug washout (shaded area) reveals no significant difference (paired t-test; p > 0.05). (B) 10 min pretreatment of the nonbiased M_1_ PAM VU0453595 (10 μM), followed by a 10 min co-application of VU0453595 + 10 μM CCh, produced a significant LTD (81.5 ± 4.70%, n/N= 14/11; paired t-test p = 0.01). Neither (C) VU0405652 (10 μM) (93.5 ± 3.28%, n/N= 9/8; paired t-test; p > 0.05) nor (D) VU0405645 (10 μM) (91.9 ± 4.67%, n/N= 7/7; paired t-test; p > 0.05) induced a significant LTD. (E) Summary of the last 5 min of the recordings from the time course experiments (& = p < 0.05, paired t-test). (F) A high concentration of CCh (50 μM, black) induces a LTD of fEPSPs (70.0 ± 7.78%, n/N= 9/7; paired t-test p = 0.003) whereas VU0405645 (10 µM, gray) blocked CCh (50 µM) induced LTD of fEPSPs (101 ± 8.59%, n/N= 11/8; paired t-test; p > 0.05). (G) Summary of the last 5 min of the recordings from the time course experiments reveals a significant difference between CCh (50 µM) alone and CCh (50 µM) in the presence of VU0405645 (10 µM) (Student’s t-test; p = 0.017). Insets in panel A, B, C, D and F show representative fEPSP traces for each condition for baseline (red trace) and 50 min after CCh washout (black trace), and scale bars denote 0.5 mV and 5 ms. Data are expressed as mean ± SEM. * p < 0.05, **p < 0.01.

Theoretically, M_1_ PAMs that confer this form of biased M_1_ signaling stabilize a conformation of M_1_ that favors activation of signaling by PLC and not PLD (*43–46*). Based on this, if these PAMs confer true bias to M_1_ signaling, they should inhibit PLD-mediated responses. Thus, we tested the hypothesis that PAMs that bias M_1_ signaling away from PLD would therefore block the LTD normally induced by a maximal concentration of CCh (Fig. 3F). In agreement with our hypothesis, VU0405645 blocked CCh-induced LTD (Fig. 3F, G). Collectively, these findings demonstrate a novel role of PLD in this cortical M_1_-LTD and that biased M_1_ PAMs not only fail to potentiate a submaximal concentration of CCh-LTD, but can also actively block a maximal concentration of CCh-LTD.

### PLD is not necessary for the M_1_-dependent increase of layer V sEPSC in the mPFC

In light of these findings, we next set out to determine whether PLD is important in other M_1_-dependent functions in the CNS. Previously, we reported that M_1_ activation increases the frequency of spontaneous excitatory postsynaptic currents (sEPSC) in mPFC layer V pyramidal neurons (*15, 19, 30*). In agreement with these previous findings, bath application of a maximal concentration of CCh induces a robust increase in sEPSC frequency in layer V pyramidal neurons (Fig. 4A). In contrast to M_1_-dependent LTD, the effect of CCh on sEPSCs was unchanged by pretreatment and co-application of the dual PLD inhibitor ML299 (Fig. 4B). Quantification of the peak CCh effect on sEPSC frequency indicates no statistically significant difference between CCh alone and CCh in the presence of ML299 (Fig. 4C). These data suggest that PLD is not necessary for this M_1_-dependent increase of sEPSC frequency in mPFC layer V pyramidal neurons.

**Figure 4:**
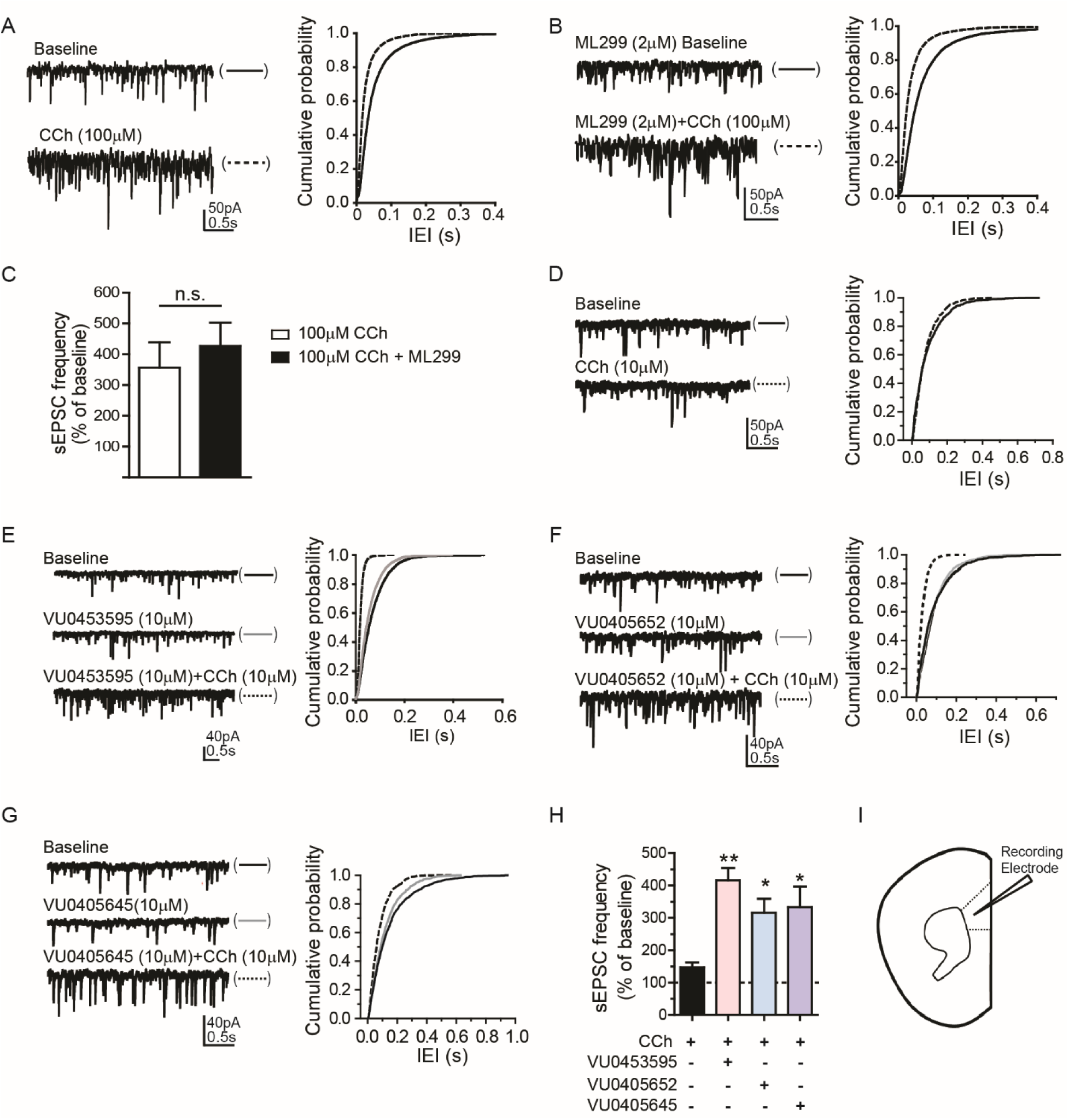
PLD is not required for M_1_-dependent increase in sEPSC frequency in mPFC layer V pyramidal neurons and both biased and nonbiased M_1_ PAMs can potentiate this response. (A) Sample traces (left) and the cumulative probability of interevent interval (IEI) (right) of sEPSCs in baseline) and during application of 100 µM CCh as indicted for a typical cell. (B) Sample traces (left) and the IEI cumulative probability (right) of sEPSCs in baseline with PLD_1,2_ inhibitor ML299 (2 µM) and during application of a combination of ML299 and CCh (100 µM) for a typical cell. (C) Summary bar graph demonstrates no significant difference in increases in sEPSC frequency between CCh alone (357.0 ± 81.6%, n/N = 7/3 cells/animals) and CCh in the presence of ML299 (427.0 ± 76.5%, n/N = 8/3) (Student’s t-test; p > 0.05). (D) Sample traces (left) and IEI cumulative probability (right) of sEPSCs in baseline and during application of 10 µM CCh from a typical cell. (E-G) Sample traces (left) and IEI cumulative probability (right) of sEPSCs in baseline, during application of a PAM) and the PAM with CCh as indicated for typical cells are shown. (H) Quantification of the peak effect on sEPSC frequency indicates a statistically significant difference between 10 µM CCh alone (147 ± 15.4%, n/N = 7/3), CCh with VU0453595 (10 μM) (416 ± 38.2%, n/N = 8), CCh with VU0405652 (10 μM) (316 ± 43.3%, n/N = 10), and CCh with VU0405645 (10 μM) (332.4 ± 63.7%, n/N = 11). One-way ANOVA F_3,35_ = 5.77; p = 0.0026, with a post-hoc Dunnett’s test using CCh alone as the control group, * p < 0.05, ** p < 0.01. Data are expressed as mean ± SEM. (I) Schematic of whole-cell recordings from mPFC layer V pyramidal neurons (regular spiking cells) clamped at −70 mV.

Next, we sought to compare the two biased M_1_ PAMs, VU0405652 and VU0405645, to our prototypical M_1_ PAM, VU0453595, in their ability to potentiate a submaximal concentration of CCh-induced increases in mPFC layer V pyramidal neuron sEPSC frequency. As expected, bath application of a submaximal concentration of CCh did not induce a significant change in sEPSC frequency (Fig. 4D). Similar to the M_1_ PAM BQCA (*15*), the non-biased M_1_ PAM VU0453595 induces a robust potentiation of the effect of a submaximal concentration of CCh on sEPSC frequency (Fig. 4E). Consistent with the studies with PLD inhibitors, both VU0405652 (Fig. 4F) and VU0405645 (Fig. 4G) potentiate agonist-induced increases in sEPSC frequency. Quantification of the peak effect on sEPSC frequency indicates a statistically significant difference between CCh alone and all three M_1_ PAMs (Fig. 4H). Therefore, both biased and non-biased M_1_ PAMs function similarly in their ability to potentiate a submaximal concentration of agonist-induced increases in mPFC layer V pyramidal neuron sEPSC frequency (Fig. 4I).

### PLD is not necessary for M_1_ effects on excitability of striatal SPNs

The M_1_ receptor is also highly expressed in the striatum (*47*), and we have shown that M_1_ activation in spiny projection neurons (SPN) in the dorsal lateral striatum leads to a robust increase in SPN excitability that can be blocked by a selective M_1_ antagonist (*48, 49*). Therefore, we set out to determine whether PLD is required for this M_1_-dependent response. As expected, bath application of CCh induces a robust increase in dorsal lateral striatum SPN excitability (Fig. 5A). In the presence of the dual PLD inhibitor ML299, CCh is still able to induce a marked increase in SPN excitability compared to baseline (Fig. 5B). Quantification of the CCh-induced increase in SPN excitability showed no significant difference between the change in number of spikes per pulse between control (DMSO) and ML299 groups (Fig. 5C). Therefore, similar to the sEPSC findings, PLD is not necessary for M_1_-dependent increases in dorsal lateral SPN excitability.

**Figure 5:**
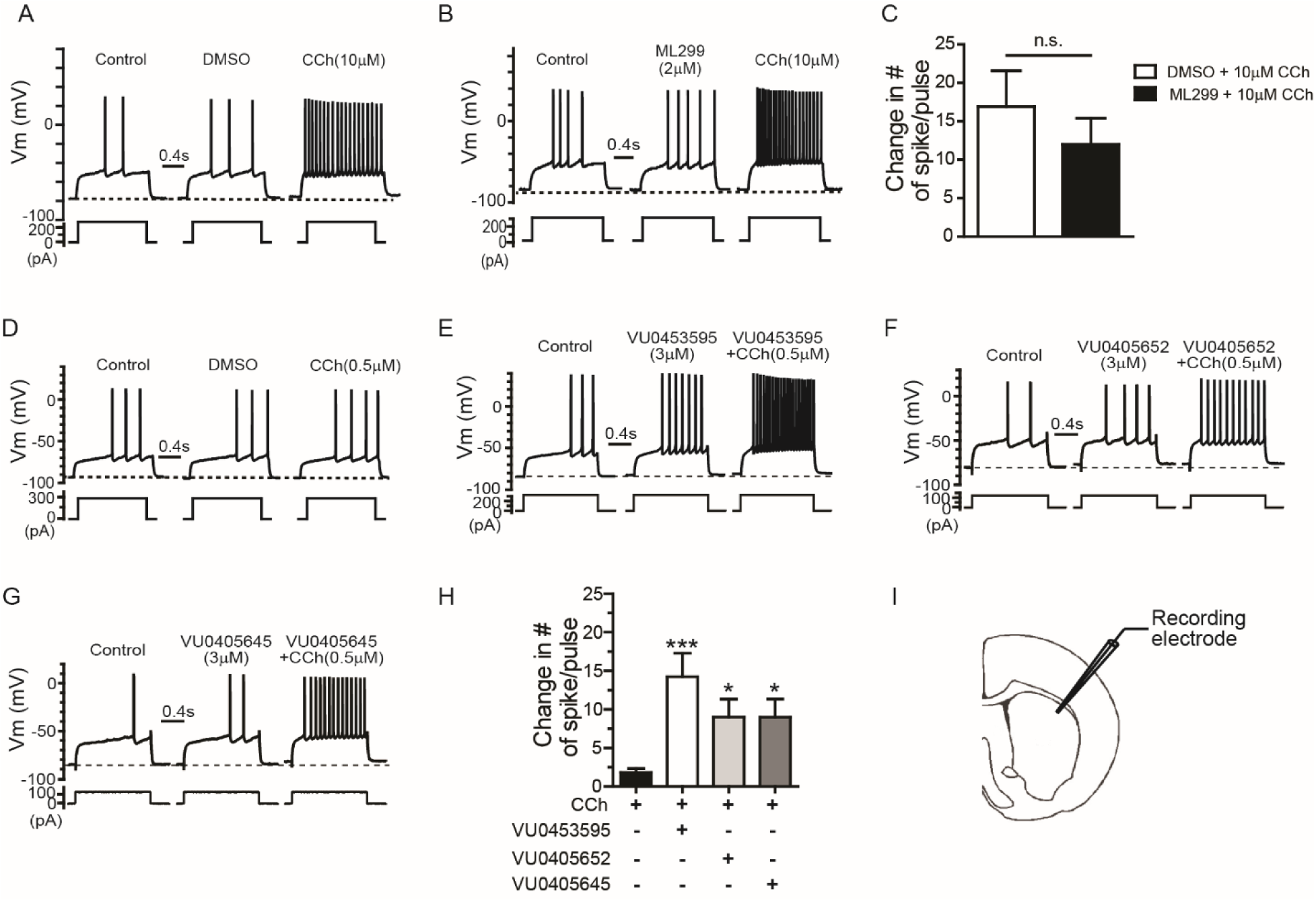
PLD is not necessary for M_1_ effects on excitability of striatal SPNs and both biased and nonbiased M_1_ PAMs can potentiate this response. (A-B) Sample traces of membrane potential responses to a depolarization current step from an SPN during baseline, in the presence of DMSO or ML299 (2 µM), then carbachol (10 µM). (C) Bar graph summarizing the changes in the number of spikes per pulse after CCh application in presence of ML299 (12.0 ± 3.38, n/N = 6/5 cells/animal) or DMSO (16.9 ± 4.67, n/N = 5/5) shows no significant difference between groups (Student’s t-test; p > 0.05). (D-G) Sample traces of membrane potential responses to a depolarization current step from an SPN during baseline, in the presence of M_1_ PAM (10 µM) or DMSO, then M_1_ PAM/DMSO + CCh (0.5 µM). (H) Bar graph summarizing the change in the number of spikes per pulse after CCh (0.5 µM) application in presence of DMSO (1.83 ± 0.49, n/N = 9/7), VU0453595 (14.2 ± 3.05, n/N = 6/6), VU0405652 (9.02 ± 2.31, n/N = 7/6) and VU0405645 (9.00 ± 2.37, n/N = 6/5) shows significant increase in the change in number of spikes/pulse compared to DMSO controls (one-way ANOVA F_3,24_ = 6; p = 0.0017, with a post-hoc Dunnett’s test using CCh + DMSO as the control group, * p < 0.05, *** p < 0.001). Data are expressed as mean ± SEM. (I) Schematic of whole-cell recordings from SPN neurons under current clamp conditions performed in the DLS.

The finding that PLD is not involved in M_1_ regulation of SPN excitability suggests that biased M_1_ PAMs that selectively potentiate coupling to PLC and do not potentiate PLD activity would function similarly to non-biased M_1_ PAMs in their ability to potentiate responses to a low concentration of CCh on SPN excitability. In agreement with our previous findings (*49*), a submaximal concentration of CCh induces a minimal increase in SPN excitability (Fig. 5D) that can be robustly potentiated by the prototypical M_1_ PAM VU0453595 (Fig. 5E). As expected, both VU0405652 (Fig. 5F) and VU0405645 (Fig. 5G) potentiate a submaximal concentration of CCh-induced increase of SPN excitability. The maximal increase in the number of spike discharges during agonist application was significantly higher in the presence of each of the three M_1_ PAMs compared to the DMSO control condition (Fig. 5H-I). Taken together, these results demonstrate that biased and non-biased M_1_ PAMs function similarly in their ability to potentiate M_1_-dependent CNS responses that are PLD-independent.

## Discussion

A large body of clinical and preclinical research suggests that enhancing mAChR signaling can be efficacious in the treatment of the cognitive symptoms associated with Alzheimer’s disease and schizophrenia (*7, 8, 50, 51*). While multiple mAChR subtypes are involved in regulation of cognitive function, the majority of preclinical studies point to a dominant role of M_1_ and suggest that selective modulation of the M_1_ receptor may provide a therapeutic potential for the treatment of these devastating cognitive symptoms (*14, 15, 17–19, 30, 52–58*). However, M_1_ regulates CNS function by actions on multiple signaling pathways and M_1_ PAMs can display a strikingly diverse range of pharmacological properties, including signal bias. At present, little is known about the specific signaling pathways involved in the different physiological effects of M_1_ activation and how signal bias can affect PAM modulation of M_1_ actions in identified brain circuits. The present studies provide a fundamental advance in our understanding of at least one novel mechanism by which M_1_ activation leads to plasticity changes within a key cortical structure in the CNS. Specifically, we found that a previously described M_1_-dependent LTD in the cortex is dependent on activation of PLD. Furthermore, we identified novel M_1_ PAMs that selectively enhance M_1_ coupling to PLC but not PLD, and found that these biased M_1_ PAMs fail to potentiate this form of M_1_-dependent LTD. Finally, these biased M_1_ PAMs actively block the ability of mAChR agonists to induce this PLD-dependent LTD, consistent with the hypothesis that these PAMs stabilize a conformation of M_1_ that favors activation of PLC over PLD and thereby bias M_1_ signaling in favor of PLC-mediated responses. Importantly, not all M_1_-dependent responses are PLD-dependent and biased M_1_ PAMs function similarly to non-biased M_1_ PAMs in M_1_ signaling that is PLD-independent.

While the ability of M_1_ and other GPCRs to activate PLD is well established (*59*), little is known about the physiological roles of PLD in regulating CNS function. This has largely been due to the lack of selective inhibitors and other tools that allow systematic studies of PLD-mediated responses. However, the recent discovery of the highly selective PLD inhibitors used here (*35–37*), along with generation of PLD KO mice and biased M_1_ PAMs reported in the present studies, provided an unprecedented opportunity to determine the roles of PLD in mediating specific responses to M_1_ activation. With the availability of these new tools, these studies provide the first clear example of a specific physiological role of PLD in mediating a response to GPCR activation in the CNS and reveal a novel role for PLD in the induction of major form of synaptic plasticity in an identified brain circuit. Furthermore, these novel PLD inhibitors include selective inhibitors of PLD_1_ and PLD_2,_ the major isoforms of PLD expressed in the CNS. These isoform-selective inhibitors, along with PLD_1_ KO mice, reveal a critical role of PLD_1_ as the PLD isoform involved in mediating this response to M_1_ activation.

M_1_-dependent LTD in the mPFC has been extensively studied and has been postulated to play a critical role in regulating specific inputs to the mPFC from the hippocampus and other extrinsic afferents (*60*). Cholinergic regulation of these inputs is thought to be important for the regulation of multiple aspects of mPFC function and previous studies suggest that M_1_ expression and signaling in the mPFC can be impaired in some pathological states that could be relevant for schizophrenia and Alzheimer’s disease (*3, 14, 30, 61–67*). However, very few studies have focused on understanding the cellular mechanisms underlying M_1_-dependent LTD in the PFC. While the current studies identify PLD_1_ being critically important in M_1_-dependent cortical synaptic plasticity, the detailed molecular mechanism by which M_1_ signals through PLD to induce synaptic plasticity changes in the cortex remains unknown. Rigorous molecular and biochemical studies to elucidate this signaling pathway are necessary to fully understand the signaling cascade responsible for M_1_-dependent LTD.

The finding that PLD_1_ is important for this form of synaptic plasticity, coupled with the finding that biased and non-biased M_1_ PAMs have functionally distinct effects on this response, raises the possibility that different PAMs could have unique profiles in regulating cognitive function or other in vivo responses. It is possible that biased versus non-biased M_1_ PAMs could induce dramatically different effects on specific behavioral responses, as is the case for biased and non-biased PAMs of the mGlu_5_ subtype of metabotropic glutamate (mGlu) receptor (*68, 69*). Unfortunately, the currently available biased M_1_ PAMs used in the present studies do not have appropriate properties to allow their use in in vivo behavioral studies. However, in future studies it may be possible to optimize biased M_1_ PAMs that can be used to systematically evaluate the roles of PLD in specific behavioral responses that are dependent on M_1_ activation. Extensive medicinal chemistry efforts are needed to develop biased M_1_ PAMs that have favorable physical and pharmacokinetic properties suitable for systemic administration with high CNS penetrance in order to test whether systemically administered biased M_1_ PAMs display functional differences in their ability to reverse cognitive deficits in preclinical animals relevant for AD and schizophrenia.

Finally, in future studies, it will also be important to develop an understanding of the precise molecular mechanisms involved in conferring bias for some M_1_ PAMs. While there are multiple examples of allosteric modulators of GPCRs inducing biased signaling, little is known about the structural basis of biased versus non-biased signaling. Previous studies reveal multiple allosteric binding sites for some GPCRs, which could contribute to different responses to distinct classes of allosteric modulators (*70–72*). However, recent studies suggest that differences in M_1_ PAM functionality may not be due to binding to different allosteric binding pockets, but that binding of PAMs to a single allosteric site may stabilize different receptor conformational states (*73, 74*). Understanding how allosteric modulators of GPCRs induce their effects will help facilitate the rational design of the next generation of PAMs and NAMs.

## Materials and Methods

### Cell line and Calcium Mobilization Assay

Briefly, M_1_-CHO cells were plated in black-walled, clear-bottomed 384 well plates (Greiner Bio-One, Monroe, NC) the day before assay. The next day, cells were washed with assay buffer (Hank’s balanced salt solution, 20 mM HEPES, 4.16 mM sodium bicarbonate, and 2.5 mM probenecid) and immediately incubated with 20 μL of 1.15 μM Fluo-4-acetomethoxyester (Fluo-4 AM) dye solution prepared in assay buffer for 45 min at 37 °C. M_1_ PAMs were serial diluted (1:3) in DMSO for 10-point concentration−response curves (CRC), and further diluted in assay buffer at starting final concentration 30μM using Echo liquid handler (Labcyte, Sunnyvale CA). After removing dye, cells were washed with assay buffer and immediately calcium flux was measured using the Functional Drug Screening System (FDSS7000, Hamamatsu, Japan) The serially diluted compounds or DMSO vehicle were added to cells for 2.5 min and then an EC_20_ concentration of acetylcholine (ACh) was added and incubated for 2 min. EC_max_ concentration was also added to cells that were incubated with DMSO vehicle to ensure the EC_20_ calcium response. To determine the potency and efficacy of the agonist and PAM, data were analyzed to generate a concentration-response curve using a four-point logistical equation in GraphPad Prism 5.0 (GraphPad Software, Inc., La Jolla, CA).

### PLD activity assay

Methods were adapted from Walker SJ & Brown HA, 2004. Briefly, CHO cells stably transfected with the human M_1_ muscarinic acetylcholine receptor were grown on growth media consisted of Ham’s F-12 Nutrient Mix (ThermoFisher #11765), 10% FBS, 20 mM HEPES, 1X Antibiotic/Antimycotic, 500 µM G418. The cells were then plated on 6-well plates for a total of approximately 0.7×10^6^ cells / 2mL / well. Plating media consisted of growth media without FBS or G418. The following day, plating media was aspirated off and labeling media was prepared by adding ^3^H-palmitic acid (5 µCi/ µL) supplemented with 2.08 µg/µL Phosphoethanolamine (PE stock, 25 mg/mL in CHCl_3_) to serum free media supplemented with bovine serum albumin. Each well contained 1mL media with 10 to 30 µCi [^3^H]-palmitic acid. Labeling was allowed to occur in a 37 °C incubator overnight.

The next morning, the plating media was carefully aspirated off and the cells were treated for 5 min with DMSO or M_1_ PAM then 30 min in the presence of 0.3% 1-butanol in serum-free assay media (1 mL media/well) or no butanol serum-free assay media as a negative control and the plates were incubated at 37 °C. ^3^H labeling efficiency was measured by subtracting the post-labeling medium from the pre-labeling medium. All pharmacological agent stocks were used at 500 or 1000-fold higher than the final concentration. Immediately after the incubation, 600 µL ice cold acidified methanol (1:1 ratio of 0.1 N HCl to Methanol) was added and the cells were scraped off using a cell scraper and transferred to a 1.5 mL Eppendorf tube. 300 µL room-temperature CHCl_3_ was then added and the sample which was then vortexed vigorously for approximately 20 seconds. The samples were then spun at 16,000 g for 5 min to separate phases. The bottom lipid phase was removed carefully to ensure no other phases were carried over and transferred to a new 1.5 mL Eppendorf tube. The samples were then dried under N_2_ gas until all liquid was evaporated. The lipids were then resuspended in 25 µL CHCl_3_ and immediately spotted onto the TLC plate (Sorbtech, Norcross GA; Cat#2315126C). Non-radioactive lipid standards such as p-Butanol and phosphatidic acid were also spotted on the TLC plate. The TLC tank was prepared by placing chromatography paper 7-inch (H) x 22.5-inch (W) so that it covers approximately 75% of the tank’s height. The mobile phase was then added (10 CHCl_3_: 2 Methanol: 2 Acetic Acid: 4 Acetone: 1 H_2_0) and allowed to equilibrate for 1 hour before the TLC plate was added and run for 1.5 to 2 hours. The plate was then removed from the tank and allowed to completely dry before imaging using autoradiography film in conjunction with an intensifying screen (BioMax Transcreen LE, Carestream Health) and placed in a −80 °C freezer for 3-5 days. The film was then processed after exposure and quantified using Chemdoc (Biorad).

### Animals

All animal studies were approved by the Vanderbilt University Medical Center Institutional Animal Care and Use Committee and were conducted in accordance with the National Institutes of Health Guide for the Care and Use of Laboratory Animals. 6-10 week old male and female C57BL6/J mice (Jackson laboratories) as well as both male and female PLD1 knockout (KO) mice (obtained from the trans-NIH Knock-Out Mouse Project Repository, www.komp.org) maintained on a C57BL6/J background were used in electrophysiology studies. Mice were group housed 4–5 per cage, maintained on a 12 h light/dark cycle, and food and water were provided ad libitum.

### Whole cell electrophysiology

Mice were anesthetized with isoflurane then transcardially perfused with ice-cold cutting solution (in mM: 230 sucrose, 2.5 KCl, 8 MgSO_4_, 0.5 CaCl_2_, 1.25 NaH_2_PO_4_, 10 D-glucose, 26 NaHCO_3_), and the brains were removed then submerged in ice-cold cutting solution. Coronal slices containing either dorsal striatum or prelimbic prefrontal cortex (PFC) were cut at 250 µm or 300 µm, respectively, and were transferred to a holding chamber containing NMDG-HEPES recovery solution (in mM: 93 NMDG, 2.5 KCl, 1.2 NaH_2_PO_4_, 30 NaHCO_3_, 20 HEPES, 25 D-glucose, 5 sodium ascorbate, 2 thiourea, 3 sodium pyruvate, 10 MgSO_4_, 0.5 CaCl_2_, 12 N-acetyl-L-cysteine, pH 7.35, <310 mOsm) for 8-10 minutes at 32 °C. Slices were then transferred to a room temperature holding chamber for 1.0 hour containing ACSF (in mM: 126 NaCl, 1.25 NaH_2_PO_4_, 2.5 KCl, 10 D-glucose, 26 NaHCO_3_, 2 CaCl_2_, 1 MgSO_4_) supplemented with 600-µM sodium ascorbate for slice viability. All buffers were continuously bubbled with 95% O_2_/5% CO_2_. Subsequently, slices were transferred to a 30-31 °C submersion recording chamber (Warner Instruments) where they were perfused with ACSF at a rate of 2 mL/min. Recording pipettes were constructed from thin-walled borosilicate capillary glass tubing (I.D.=1.17 mm, O.D. 1.50 mm; Warner Instruments, Hamden, CT), pulled with a horizontal pipette puller (P-97 Sutter Instrument Co., Novato, CA) to a resistance of 4-6 MΩ when filled with potassium-based internal solution: (mM) 125 K-gluconate, 4 NaCl, 10 HEPES, 4 MgATP, 0.3 NaGTP, 10 Tris-phosphocreatine.

For the PFC recordings, pyramidal neurons were visualized based on morphology with a 40X water-immersion lens with oblique illumination coupled with an Olympus BX50WI upright microscope (Olympus). After a stable gigaohm seal was formed, light suction was applied to break through the cell membrane and achieve whole-cell access. The access resistance was checked at the beginning and the end of each experiment and neurons with an access resistance of neurons greater than 30 mOhm were not used for analysis. Pyramidal neurons were further identified by their regular spiking pattern following depolarizing current injections induced by a series of 500 ms current steps (−150pA to +100 pA) incremented in +25 pA performed in current clamp mode. Spontaneous EPSCs were recorded at a holding potential of −70 mV (the reversal potential for GABA_A_ channels) and the junction potential was not compensated. The voltage clamp signal was low pass filtered at 5 kHz and digitized at 10 kHz using a Digidata 1322A and acquired using Axon MultiClamp 700B (Molecular Devices, Sunnyvale, CA) and controlled by pClamp 9.2 and Clampex 10.6.2 running on a Dell PC. After a stable baseline was recorded for 5-10 min, test compounds were diluted to the appropriate concentrations in DMSO (<0.1% final) in ACSF and applied to the bath using a peristaltic pump perfusion system. Cumulative probability plots of inter-event-intervals (IEI) were constructed using 2 min episodes of baseline and peak effect during drug application. All sEPSC analyses were performed using MiniAnalysis (Synaptosoft Inc., Decatur, GA) or Clampfit 10.2 (Molecular Devices).

For striatal spiny projection neurons (SPN) recordings the change in excitability of MSN was assessed in current clamp mode by monitoring the change in the number of spike discharges in response to a near rheobase depolarization current step (1.5 s). The access resistance was checked at the beginning and the end of each experiment, which were compensated using “bridge balance”. The change in spike number was calculated by averaging the number of spikes during the baseline subtracted from the peak drug-effect (60 seconds). Offline data analysis to calculate change in SPN excitability was performed using Clampfit 10.2 (Molecular Devices).

### Extracellular field electrophysiology

400 µm coronal slices containing prelimbic prefrontal cortex were obtained as described above. Recording pipettes were constructed from thin-walled borosilicate capillary glass tubing (I.D.=1.17 mm, O.D. 1.50 mm; Warner Instruments, Hamden, CT), pulled with a horizontal pipette puller (P-97 Sutter Instrument Co., Novato, CA) to a resistance of 1-3MΩ when filled with ACSF. Field excitatory postsynaptic potentials (fEPSPs) were recorded from layer V of the prelimbic cortex and evoked electrically by a concentric bipolar stimulating electrode (200 µs duration, 0.05 Hz; inter-pulse interval of 50 ms) in the superficial layers II-III. Layer II/III was visualized using a Olympus BX50WI upright microscope (Olympus, Lake Success, NY) microscope according to landmarks illustrated in the Allen mouse brain atlas (*75*) and the recording electrode was placed laterally approximately 200 µM away from layer 2/3 into layer V so that the recording and stimulating electrodes were parallel to each other. Input-output curves were generated to determine the stimulus intensity that produced approximately 70% of the maximum fEPSP slope before each experiment, which was then used as the baseline stimulation. Data were digitized using a Multiclamp 700B, using a sampling rate of 20,000kHz and were filtered at 0.5kHz, with a Digidata 1322A, pClamp 9.2 and Clampex 10.6.2 software (Molecular Devices) running on a Dell PC (Round Rock, TX). All test compounds, with the exception of CCh (Tocris Bioscience, Bristol, UK) which was diluted in H_2_O, were diluted to the appropriate concentrations in DMSO (<0.1% final) in ACSF and applied to the bath using a peristaltic pump perfusion system. Offline data analysis to calculate fEPSP slope was performed using Clampfit 10.2 (Molecular Devices).

### Statistics

Two-tailed Student’s t-test and one-way ANOVA with Bonferonni post-tests were used as appropriate. Changes in fEPSP slope before and during drug add (peak effect) was compared using a paired t-test. For all statistical comparisons, the critical p-value was considered to be 0.05. The numbers of animals to be used for each experiment outlined within the study were determined using a power calculation statistical analysis using the Power and Sample Size Calculation software program available at Vanderbilt University (Dupont and Plummer, PS Controlled Clinical Trials. 18:274 1997). Animal numbers are based on a power calculation using standard errors from published studies and previous experience to detect >20% difference for each outlined experiment with an 80% power (alpha = 0.05, power = 80%, delta = 0.2, sigma = 0.18).

## Acknowledgments

We would like to thank the Dan Marino foundation for their generous support of the Clinical Neuroscience Scholars program at Vanderbilt University, to thank Weimin Peng for excellent assistance in maintaining colonies and genotyping of transgenic mice.

## Author Contributions

SPM, ZX, CAD, CMD, CWL and PJC designed experiments; SPM, ZX, CAD, JM, XL, SF, and JMR performed experiments. SPM and PJC wrote the manuscript with input from all the authors.

